# Salivary corticosterone reflects plasmatic levels in a wild seabird

**DOI:** 10.1101/2023.06.16.545258

**Authors:** Jeffrey Carbillet, Lauri Saks, Tuul Sepp

**Affiliations:** Institute of Ecology and Earth Sciences, University of Tartu, Tartu, 51014, Estonia; Estonian Marine Institute, University of Tartu, Mäealuse 14, 12618 Tallinn, Harju County, Estonia

**Keywords:** free-ranging birds, colonial seabird, Common gull, stress hormones, methodological tool

## Abstract

Wild animals have been increasingly exposed to a wide range of stressors, mainly due to the intensification of human activities and habitat modifications. Consequently, new tools in order to assess the physiological and health status of wild animals have been developed. In particular, glucocorticoids have received a special attention. Primarily metabolic hormones, they are also used to evaluate the stress level of organisms. While historically measured in blood samples, new less-invasive methods have been recently developed to measure glucocorticoids in matrices such as faeces, hairs/feathers, or saliva. To date, measurements in saliva are still in their infancy despite the numerous advantages of the matrix: non-invasive, reflects the biologically active portion of glucocorticoids, allows to measure both baseline and stress-induced levels. In addition, most studies using saliva have been performed on domestic and captive animals, and recent development in wild animals have focused on mammals. Here, we show, for the first time that saliva could also be reliably used in free-ranging birds, as glucocorticoid levels in saliva strongly correlated with plasma levels. This promising result opens new avenues for a non-invasive sampling method to assess health status of wild birds in conservation biology and ecology.

## Introduction

Due to the intensification of anthropogenic activities [1], increased occurrences in adverse weather events, and modifications in land use [2], wild animals have been facing an increased exposure to different kinds of stressors, acting as strong selective pressures and causing loss of biodiversity [3,4]. Monitoring health status estimates to predict and prevent population declines has therefore become a principal aim in conservation biology and ecology [5].

In vertebrates, the main coping mechanism to stressful situations is the stress response, which aims at neutralising the effects of stressors and restoring homeostasis [6, 7]. The most studied physiological stress response in vertebrates is the activation of the neuroendocrine system which results in the secretion of glucocorticoids such as cortisol and corticosterone into the blood stream. Primarily metabolic hormones ensuring the regulation of an individual’s energy balance, glucocorticoid levels increase when facing stressors [8,9]. When these stressors are persistent or regularly repeated, glucocorticoid secretion remains chronically elevated, and several negative consequences are observed on physiology, which may affect fitness and ultimately population dynamics [7,10].

To evaluate the health status of organisms and their ability to efficiently respond to environmental perturbations, glucocorticoid levels have been extensively measured in the blood of both domestic and wild animals [11,12]. Recently, new less-invasive matrices such as feathers, urine, faeces or saliva have been used, and several review articles have compared the advantages and drawbacks of each matrix [11,13,14]. Among them, saliva offers many advantages in addition to its non-invasiveness. First, it only reflects the unbound, or free, glucocorticoids from the blood that corresponds to the biologically active fraction of the hormones [13,15]. Second, while blood samples need to be collected under 2 to 3 minutes after the animal is captured to reflect baseline glucocorticoid levels [16], an increase in the hormone level is not detectable under 20 to 30 minutes in salivary samples [13,17], meaning that it is possible to measure both baseline and stress-induced levels quite easily. Finally, glucocorticoids have been shown to be very stable in saliva, thus allowing to store samples at room temperature for a few days before being frozen [13,18], while blood, urinary, or faecal samples need to be frozen as soon as possible [13].

Despite these numerous advantages, studies measuring glucocorticoids in saliva remain scarce to date, and most of them have been conducted on domestic [19,20,21], or captive animals [22,23,24], but see [25,26] for examples on free-ranging monkeys. Similarly, the measurement of salivary glucocorticoids has almost fully been restricted to mammals [19-24,27], with only recent reports on amphibians [28,29], and only one very recent study on a captive bird species [30]. However, birds in general are particularly exposed to multiple stressors such as marine and air pollution [31,32], urbanisation and habitat loss [33], or predation and adverse climatic events [34]. In particular, seabirds have been identified as one of the world’s most endangered avian groups [35], being affected by both local stressors like habitat loss or pollution on the breeding grounds, and larger scale stressors like climate change on their long migration routes.

In this context, we used the Common gull (*Larus canus*) as a model species to evaluate if salivary glucocorticoids could be used to reflect plasma corticosterone levels in a free-ranging bird. We predicted that salivary corticosterone would be positively correlated with total plasma corticosterone, and could therefore be considered as a less-invasive alternative for conservation biologists and ecologists for assessing stress hormone levels in wild birds.

## Material and methods

### (a) Study site

This study took place in a free-living colony of Common gulls located on the Kakrarahu islet in Matsalu National Park, on the West coast of Estonia (58°46′ N, 23°26′ E). The islet is situated in northern temperate climate zone, at the eastern part of the Baltic Sea, and mainly composed of limestone gravel with patches of dense vegetation of herbaceous plants which become dominant during late May and early June. During last years, approximately 1200 pairs of common gulls are breeding on the islet.

### (b) Field data collection

As part of a long-term capture-mark-recapture program initiated in 1962, common gull nests in Kakrarahu were monitored between the 2^nd^ of May and 15^th^ of June 2022. As part of another project, and in order to reduce disturbance within the colony, a total of 24 adult incubating Common gulls (10 females and 14 males) were randomly chosen for both blood and saliva sampling. The exact age was known for 20 of these birds, and ranged from 5 to 17 years, with a mean of 10.9 ± 3.8 (SD). All of these birds were caught between the 14^th^ and 25^th^ of May, between 10 am and 7 pm, and released immediately after sampling. To minimise the post manipulation nest abandonment, all birds were caught from nests using spring traps after the tenth day of incubation, as previously described and done in the same colony [36-38].

### (c) Blood and salivary samples

Up to 50 μl of blood was collected in 200-μl microvette tubes with EDTA as an anticoagulant from the brachial vein using blood lancets. Almost all blood samples were collected under 3 minutes after trapping (1.40 to 3.10 minutes, with a mean of 2.27 ± 0.43 minutes) in order to measure baseline corticosterone levels [16]. Samples were immediately centrifuged at 6 700 x g for 5 min to separate plasma from erythrocytes, and samples were placed in a cooled and light-protected polystyrene foam box for a few hours before being stored at -80°C until analysis. Salivary samples were collected right after the blood samples by gently inserting a small cotton swab in the mouth of the gull and slowly moving it for 30 seconds to ensure that the absorbent part of the swab was saturated with saliva. The swab was then transferred in a plastic tube and placed in a cooled and light-protected box for a few hours, and then stored at -20 °C for 3 weeks before being stored at -80°C until analysis. On the day of assay, salivary samples were centrifuged at 1500 x g for 15 minutes (final volumes comprised between 5 and 25 μl).

### (d) Enzyme immunoassay (EIA)

For both blood and saliva, each sample was measured in duplicate by using a commercially available EIA kit (mouse anti-rabbit IgG, Corticosterone ELISA kit, number 501320, Cayman Chemicals) previously used and validated for different bird species [39,40]. The assay was conducted following the manufacturer recommendations and samples were reconstituted and diluted (1:50 for blood samples, and 1:25 for saliva samples) with Cayman Chemical ELISA buffer to ensure that concentrations fell within the linear portion of the standard curves. Samples were run across 2 EIA plates with inter-assay coefficients of variation of 6.32% (high concentration) and 8.96% (low concentration), and intra-assay coefficients of variation of 7.8% (range = 0.2 – 12.9%). For each sample, mean values of salivary and plasma corticosterone concentrations were used for statistical analysis.

### (e) Statistical analysis

In order to evaluate the strength of the relationship between salivary and plasma corticosterone concentrations, we performed a linear correlation test on our 24 samples. Since our sample size can be considered as relatively small, we decided to use both the frequentist and Bayesian approach in order to provide stronger support to our results. First, we performed a regular t-test of the Pearson correlation coefficient, using the cor.test function from the stats package. Second, we performed a Bayesian correlation test, using the correlationBF function from the BayesFactor package [41]. Finally, in order to compare the amount of corticosterone in saliva and blood, we performed a paired t-test, using the t.test function from the stats package. All analyses were carried out with R version 3.6.0 [42].

## Results

Salivary corticosterone levels increased with plasma levels of the hormone (Figure 1). Both the frequentist (Pearson’s correlation: r = 0.58, t = 3.34, df = 22, P < 0.003) and Bayesian (median = 0.48, pd = 99.6%, % in rope = 0%, BF = 15.80) approaches provided a similar level of strong evidence for this positive correlation. Corticosterone concentrations in saliva (1.73 ± 1.13 SD) were on average 32.3% lower (paired t = -7.54; p < 0.001) compared to concentrations in plasma samples (5.35 ± 2.82 SD).

**Figure 1.**
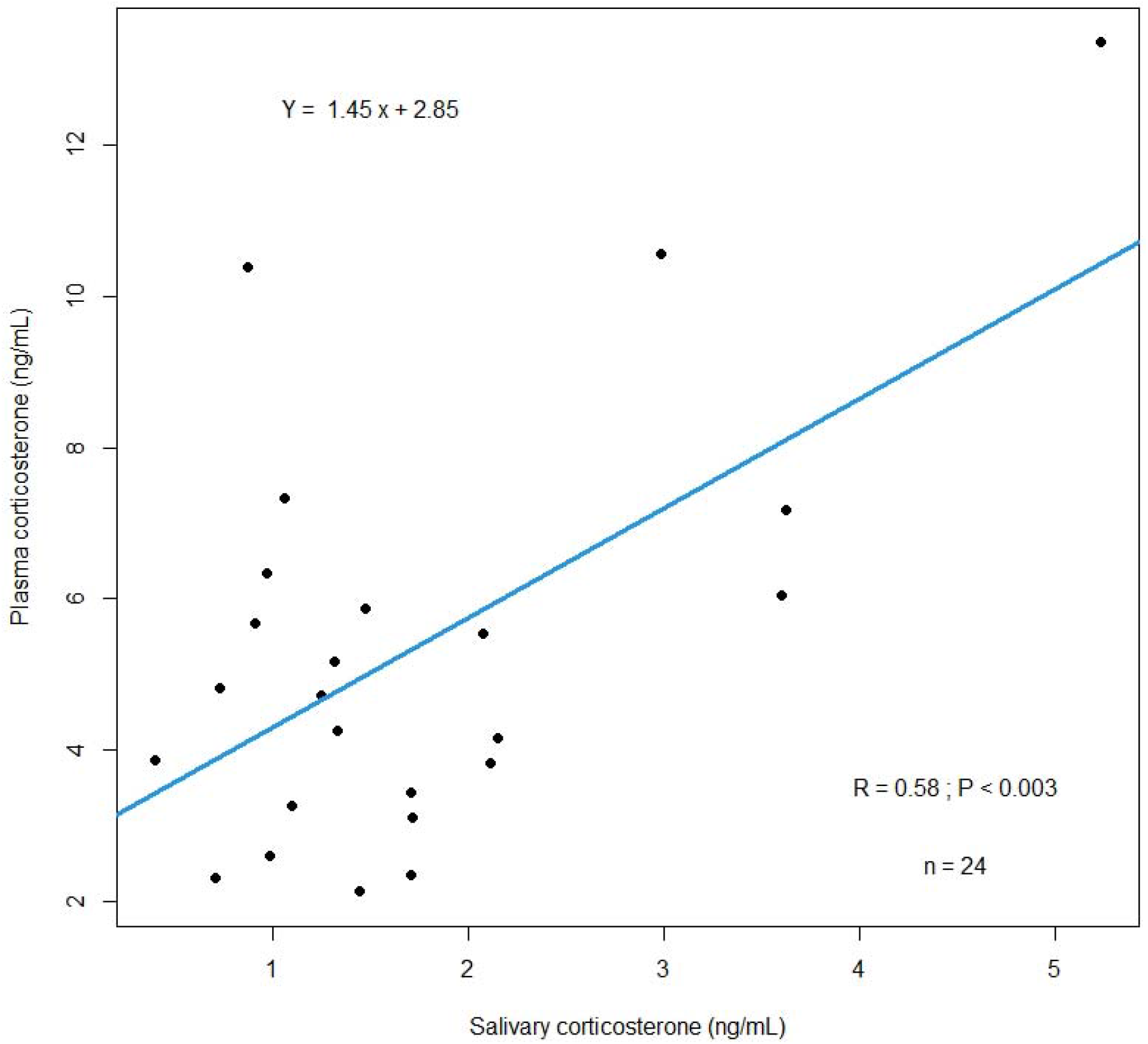
Correlation between salivary and plasma corticosterone in a free-ranging colony of common gulls (*Larus canus*). Points represent observed values.

## Discussion

Our study aimed to evaluate the potential of saliva as a non-invasive matrix to measure glucocorticoid levels in wild birds. Salivary corticosterone levels of wild Common gulls were strongly correlated to plasma levels, indicating that this matrix can be used as a less-invasive alternative for assessing stress hormone and health status in wild birds. To our knowledge, our results are the first to date to show such relationship in a wild avian model system.

When comparing salivary and plasma corticosterone concentrations, we observed that concentrations in saliva were on average only 32.3% of those found in plasma. This result is in accordance with previous studies on difference species [13,43,44] but also expected, as corticosterone in saliva reflects only the free corticosterone concentration in the blood, where corticosterone is largely bound to some proteins [13,15]. Consequently, slightly higher volumes of saliva may be required compared to plasma for analysing corticosterone concentrations. However, the fact that salivary samples reflect the free fraction of blood corticosterone makes it a premium matrix, as the free fraction of the hormone is thought to be the main biologically active one [45], and consequently the one researchers are the most interested in.

Another advantage of saliva is that it can be used to measure both baseline and stress-induced levels of corticosterone, contrary to most of the other non-invasive matrices such as faeces, hair, or urine, that reflect only baseline or long-term corticosterone changes [13,14]. Indeed, corticosterone concentrations in saliva increase around 20 to 30 minutes following a stressor [17]. This timing offers enough time to avoid measuring any manipulation-induced increase in hormonal levels. On the other side, this relatively medium time and less invasive method also offer options for repeated sampling on a short time scale to measure the intensity of corticosterone response to stressors.

While our work did not focus on performing an adrenocorticotropic hormone (ACTH) stimulation test, the only other study using salivary corticosterone in a captive bird (large-billed crows) did it and showed that the increase in corticosterone levels following ACTH injection was similar between blood and saliva, using the same commercial EIA kit than us [30]. This finding reinforces our own result on the validity and potential of saliva as a non-invasive matrix for measuring steroid hormones in free-ranging birds.

While our work focused on corticosterone, other steroid hormones and peptides could also potentially be measured in saliva of free-ranging birds, as it is already done in humans and some domestic animals [46-48]. In particular, previous studies showed that any unconjugated hormone enters saliva by free diffusion through the salivary gland cells, which makes their concentration independent on the salivary flow rate [49,50]. As such, it could be possible to non-invasively monitor several other physiological parameters in saliva, such as testosterone [51], oestradiol [52], progesterone [46], oxytocin [47], or melatonin [48] for example, providing insight on the reproductive, behavioural and overall health status of wild birds.

To conclude, to our knowledge, our study provided the first evidence in free-ranging birds that salivary collection is possible, and that salivary corticosterone levels were strongly correlated to total plasma levels. As such, our work should be seen as a steppingstone for further studies aiming to evaluate the health status of wild birds in the fields of conservation biology and ecology.

## Supporting information

electronic supplementary material 1

## Ethics

All the procedures used in this study were approved by the Ministry of Rural Affairs of the Republic of Estonia (licence no. 213, issued 24.03.2022) and was performed in accordance with relevant Estonian and European guidelines and regulations.

## Data accessibility

Datasets and code for processing and analysing data are available in the electronic supplementary material 1.

## Authors’ contributions

J.C.: conceptualization, investigation, data curation, formal analysis, funding acquisition, methodology, writing-original draft, and writing-review and editing; L.S. investigation, methodology, writing-original draft, and writing-review and editing; T.S.: investigation, funding acquisition, writing-original draft, and writing-review and editing. All authors gave final approval for publication and agreed to be held accountable for the work described in this manuscript.

## Acknowledgements

We warmly thank all the volunteers who took part to the Kakrarahu field work and helped collecting data.

