## Supplementary material for "Salivary corticosterone reflects plasmatic levels in a wild seabird": electronic supplementary material 1: script saliva BiologyLetters.R

```
###############################################################
# Saliva and plasma corticosterone in Common gulls            #
# April 2023                                                  #
# J. Carbillet                                                #
#                                                             #
###############################################################

#############
# Workspace #
#############

setwd("C:/Users/jeffreym/Documents/Donnees chevreuils")

###########
# library #
###########

library(BayesFactor)
library(bayestestR)

# Load dataset
rm(list = ls())
DataManip=read.csv("testcort.csv",header=T, dec=".")
summary(DataManip)
dim(DataManip) # 24 observations of 5 variables
attach(DataManip)

# Pearson correlation test
plot(DataManip$mean.saliva,DataManip$mean.plasma)
cor.test(DataManip$mean.saliva,DataManip$mean.plasma)

# Bayesian correlation test
result <- correlationBF(DataManip$mean.saliva,DataManip$mean.plasma,iterations = 10000)
describe_posterior(result)

# Plot
plot(DataManip$mean.saliva, DataManip$mean.plasma, pch = 16,
     ylab="Plasma corticosterone (ng/mL)", 
     xlab="Salivary corticosterone (ng/mL)")
abline(lm(DataManip$mean.plasma ~ DataManip$mean.saliva), col = 4, lwd = 3)

# Text
coef <- round(coef(lm(DataManip$mean.plasma ~ DataManip$mean.saliva)), 2)
text(1.5, 12.5,  paste("Y = ", coef[2], "x", "+", coef[1]))
text(4.5, 3.5, paste("R = 0.58",";", "P < 0.003"))
text(4.5, 2.5, paste("n = 24"))

## Paired T-test to compare salivary and plasmatic corticosterone levels
t.test(DataManip$mean.saliva, DataManip$mean.plasma, 
       paired = TRUE, alternative = "two.sided")
```
